# Mediated pleiotropy between psychiatric disorders and autoimmune disorders revealed by integrative analysis of multiple GWAS

**DOI:** 10.1101/014530

**Authors:** Qian Wang, Can Yang, Joel Gelernter, Hongyu Zhao

**Affiliations:** Program in Computational Biology and Bioinformatics, Yale University, New Haven, Connecticut, USA.; Department of Mathematics, Hong Kong Baptist University, Hong Kong SAR.; Department of Psychiatry, Yale School of Medicine, New Haven, Connecticut, USA.; VA CT Healthcare Center, West Haven, Connecticut, USA.; Department of Neurobiology, Yale School of Medicine, New Haven, Connecticut, USA.; Department of Genetics, Yale School of Medicine, West Haven, Connecticut, USA.; Department of Biostatistics, Yale School of Public Health, New Haven, Connecticut, USA; VA Cooperative Studies Program Coordinating Center, West Haven, Connecticut, USA.

**Keywords:** GWAS, Autoimmune, Psychiatric disorders, Pleiotropy, eQTL

## Abstract

Epidemiological observations and molecular-level experiments have indicated that brain disorders in the realm of psychiatry may be influenced by immune dysregulation. However, the degree of genetic overlap between immune disorders and psychiatric disorders has not been well established. We investigated this issue by integrative analysis of genome-wide association studies (GWAS) of 18 complex human traits/diseases (five psychiatric disorders, seven autoimmune disorders, and others) and multiple genomewide annotation resources (Central nervous system genes, immune-related expression-quantitative trait loci (eQTL) and DNase I hypertensive sites from 98 cell-lines). We detected pleiotropy in 24 of the 35 psychiatric-autoimmune disorder pairs, with statistical significance as strong as *p*=3.9e-285 (schizophrenia-rheumatoid arthritis). Strong enrichment (>1.4 fold) of immune-related eQTL was observed in four psychiatric disorders. Genomic regions responsible for pleiotropy between psychiatric disorders and autoimmune disorders were detected. The MHC region on chromosome 6 appears to be the most important (and it was indeed previously noted (1-3) as a confluence between schizophrenia and immune disorder risk regions), with many other regions, such as cytoband 1p13.2. We also found that most alleles shared between schizophrenia and Crohn’s disease have the *same* effect direction, with similar trend found for other disorder pairs, such as bipolar-Crohn’s disease. Our results offer a novel bird’s-eye view of the genetic relationship and demonstrate strong evidence for mediated pleiotropy between psychiatric disorders and autoimmune disorders. Our findings might open new routes for prevention and treatment strategies for these disorders based on a new appreciation of the importance of immunological mechanisms in mediating risk.

## Significance Statement

Co-occurrence of psychiatric disorders and autoimmune disorders has long been recorded while their shared genetic factors are less well explored and remain controversial. We performed comprehensive genome-level analysis on those two classes of disorders by integrating both disorder-specific genome-wide association studies (GWAS) and genomic annotations, in search of common genetic liability. Our results confirmed previously reported genetic regions affecting disease risk for both psychiatric and autoimmune disorders, and also implicated many novel shared genes and pathways. Our work offers insights on pleiotropic mechanisms and a better understanding of pathophysiology, which may lead to improved prevention and treatment strategies for these two classes of disorders via immunological mechanisms.

## Introduction

Psychiatric disorders are often associated with significant morbidity and mortality (4).

The estimated heritability for most psychiatric disorders is moderate to high (40%-80%), so genetic factors play a critical role in their etiology (5-7). In the past few years, many genome-wide association studies (GWAS) have been conducted to identify genetic risk variants that underlying psychiatric disorders (8). Despite recent progress, there still exist major challenges, and there is much yet to be discovered regarding the genetic architecture of psychiatric disorders (9).

The relationship between psychiatric disorders and autoimmune disorders has intrigued researchers for decades (**Fig. S1**). There is a moderately large body of evidence that supports a role for autoimmune dysfunction in the development of several psychiatric disorders, including early hypothesis like the “macrophage theory of depression” (10), and recent findings such as the epidemiological observation of co-occurrence of rheumatoid arthritis (RA) and depression (11, 12) and cross-disorder drug effects, for example some drug for psychiatric disorders have anti-inflammatory properties (13, 14). The genetic liability underlying these observed correlations has not been well studied, with the exception that recent GWAS have repeatedly identified association between SCZ and genetic variants at the major histocompatibility locus (MHC), which also plays an important role in the immune system (1-3). However, no strong evidence of shared liability was observed between Crohn’s disease (CD) and multiple psychiatric disorders in another study (15). In genetics, the term “pleiotropy” is commonly used to describe a one-to-many relationship between a gene or mutation and phenotypes (16). In the GWAS era, pleiotropy could explain correlations among disorders, and may also boost statistical power to detect genetic associations (15, 17-21). To date, pervasive pleiotropic effects have been discovered in autoimmune disorders (22) and in psychiatric disorders (9, 15), as separate classes.

Given the public health significance of these two classes of disorders and the treatment implications of any etiological overlap, it is important to resolve the nature of any genetic pleiotropy between them, to understand the underlying mechanisms of the pleiotropy, and to identify specific genes and pathways driving such pleiotropy. These inquiries can only now be carried out because of the large amounts of genomic data that have become available in recent years. Large consortia have been formed to study many psychiatric disorders and autoimmune disorders (15, 21, 23-30). For example, the analysis results from a well-powered GWAS of schizophrenia (1) provided strong supporting evidence for the link between schizophrenia and the immune system. Undoubtedly, the availability of high-quality “*omics*” data offers us an unprecedented opportunity to revisit the nature of the genetic connections between psychiatric disorders and immune-mediated disorders. The analysis results can deepen our understanding of the genetic architecture of complex human diseases.

Our current study takes advantage of multiple “*omics*” data resources to obtain a bird’seye view of the shared genetic components between psychiatric disorders and autoimmune disorders. To better represent those two disorder categories while taking the data availability into account, we considered five psychiatric disorders, including schizophrenia (SCZ), bipolar affective disorder (BPD), autism spectrum disorder (ASD), attention deficit-hyperactivity (ADHD), and major depressive disorder (MDD). For immune-mediated disorders, we considered two inflammatory bowel diseases (IBDs), Crohn’s disease (CD) and ulcerative colitis (UC), and five other autoimmune disorders, including multiple sclerosis (MS), psoriasis (PS), rheumatoid arthritis (RA), systemic lupus erythematosis (SLE), and insulin-dependent diabetes mellitus (T1D). For comparisons, we also included a central nervous system degenerative disease, Parkinson’s disease (PD), and five traits related to education, height, and weight.

## Results

### Pervasive pleiotropic effects between psychiatric disorders and immune system disorders

Previous studies have shown extensive shared genetic effects among many of the five psychiatric disorders studied by the Psychiatric Genomics Consortium (PGC) (15, 21) and among multiple autoimmune disorders (22), separately. Consistent with those studies, we also observed pervasive pleiotropic effects among psychiatric disorders and among immune-related disorders (Table S1). Pleiotropic effects are significant (Bonferroni-adjusted p<0.05) for all 21 pairs of autoimmune disorders, and for seven of the 10 pairs of psychiatric disorders (the exceptions being ASD-ADHD, MDD-ADHD, and MDD-ASD).

We then tested pleiotropic effects between psychiatric and autoimmune disorders. We first considered SCZ with seven immune-mediated disorders. The stratified-QQplots (**Fig. 1**, **Fig. S2**) suggest that all seven immune-mediated disorders share genetic liability components with SCZ, driving the conditional observed-expected curves substantially above the baseline.

**Figure 1.**
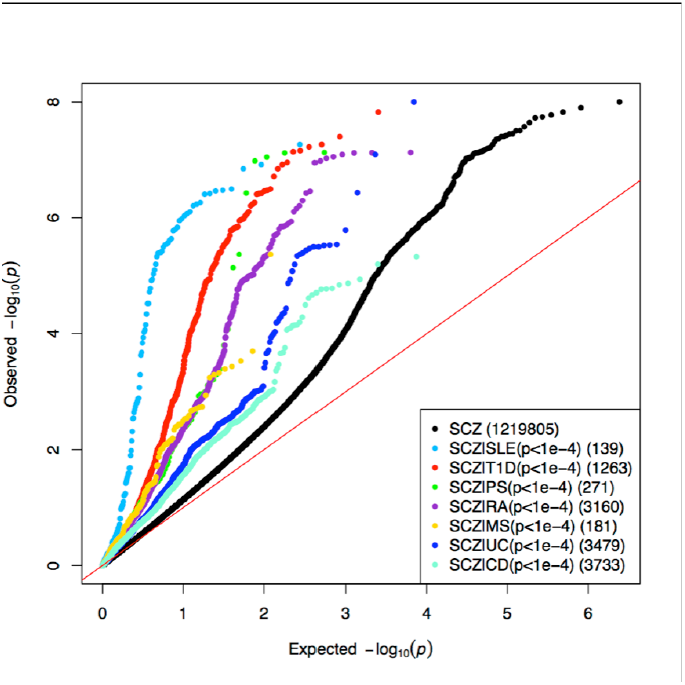
QQ plot showing pervasive pleiotropic effects between SCZ and seven immune mediated disorders. Black dots represent all 1,219,805 SCZ GWAS SNPs while the other 7 colored dots represent different subsets of SNPs selected from the corresponding autoimmune disorder GWAS whose p values were < 0.0001, with the number of SNPs in each subset shown in brackets.

QQplots, while simple and intuitive, suffer from arbitrary cutoffs, e.g. 1e-4, and do not offer statistical assessment of pleiotropy. GPA, a statistically rigorous approach recently developed by us (31), permits statistical inference regarding the genetic architecture of multiple traits (detailed in **SI**). We used GPA to test the cross-class enrichment of GWAS signals between five Psychiatric Genomics Consortium (21) (PGC) traits (SCZ, BPD, MDD, ASD, ADHD) and seven immune-mediated disorders (CD, UC, MS, PS, SLE, RA, T1D), resulting in 35 disorder pairs. Twenty-four of the 35 pairs were significant at Bonferroni-adjusted *p* value <0.05 (Table S1), indicating pervasive pleiotropic effects between psychiatric disorders and immune mediated disorders. Of the 11 pairs that were not significant, seven included ADHD, which had relatively weak GWAS signals – relatively strong GWAS signals are necessary to provide adequate information to identify pleiotropic effects; while the other four disorder pairs were ASD-SLE, BPD-MS, BPD-PS, and MDD-CD. Consistent with previous studies (32), we observed strong pleiotropy between SCZ-MS (*p*=1.3e-20), but no significant pleiotropy between BPD-MS (*p*=0.26).

For each pair of disorders, we estimated the proportion of single nucleotide polymorphisms (SNPs) associated with both disorders vs. those associated with only one disorder. **Fig. 2** shows the results among SCZ, BPD, UC and CD (**Fig. 2**). Consistent with previous studies (33, 34), most of the SCZ-associated SNPs and BPD-associated SNPs were estimated to be shared between these two disorders. Similarly, most UC-associated SNPs and CD-associated SNPs were shared between them. The proportions of SNPs shared by cross-class disorders were: SCZ-CD 0.063 (*se*=0.0021); SCZ-UC 0.053 (*se*=0.0018); BPD-CD 0.05 (*se*=0.0034); and BPD-UC 0.039 (*se*=0.0025), respectively. Proportions of shared SNPs for other disease pairs can be found in Table S1.

**Figure 2.**
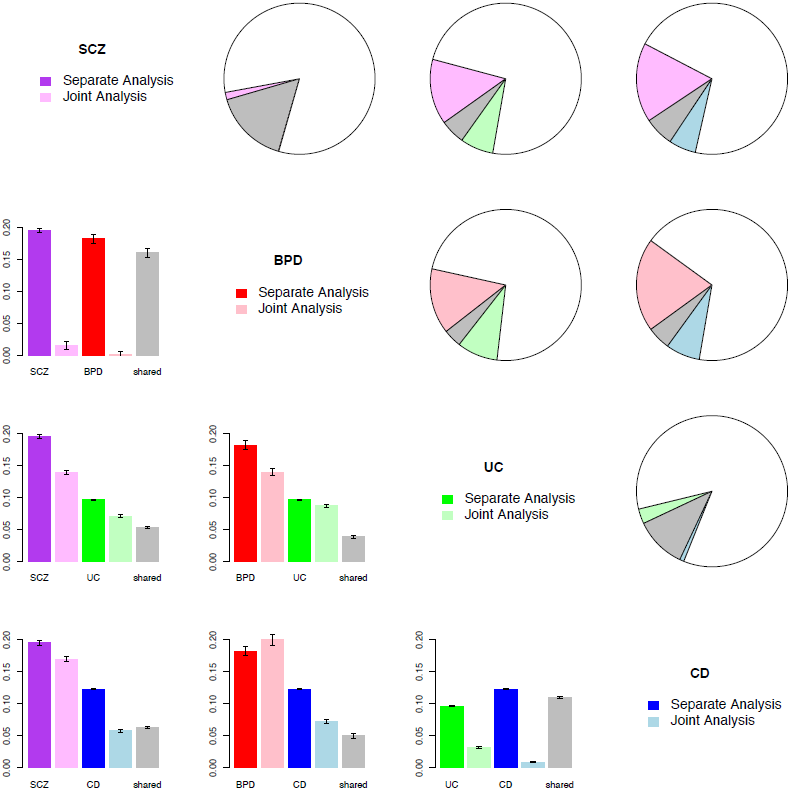
GPA results showing pleiotropic effects among SCZ, BPD, UC and CD. Purple, red, green and blue represent SCZ, BPD, UC and CD; grey represents the proportion of SNPs associated with both disorders, and white represents the proportion of SNPs associated with neither disorder. Upper triangle: pie charts show proportion of SNPs associated with only one disorder, both disorders (grey), and neither disorder (white). Lower triangle: bar plots contrasting proportions of associated SNPs for each disorder when analyzed separately (1^st^, 3^rd^ bar in deeper color), and proportion of associated SNPs when two disorders are jointly analyzed (2^nd^ and 4^th^ bar for proportion of SNPs associated with only one disorder, and 5^th^ grey bar for proportion of SNPs associated with both disorders). Error bars indicate one standard error.

### Enrichment of immune-related annotations in multiple psychiatric disorders

Having observed extensive pleiotropy between psychiatric and immune disorders, we then explored functional enrichment for the shared genes to begin to understand the biology underlying the statistical association. We used central nervous system (CNS) SNPs and immune related eQTLs (see **Materials and Methods**) to represent the functional sites relevant to the CNS and immune system, respectively, and tested their enrichments in various psychiatric disorders and immune-mediated disorders. Because 12.5% of CNS SNPs overlap immune eQTLs, we also tested enrichments excluding those overlapping SNPs.

We first tested for enrichment of CNS SNPs in all 18 traits (**Fig. 3**). As expected, all psychiatric disorders had modest enrichment for CNS SNPs (>1.3-fold, except for MDD, 1.09-fold). The enrichment effects could still be observed (and were even stronger for ADHD) with immune related eQTLs excluded. Only three autoimmune disorders (MS, PS and RA) showed modest enrichment for CNS SNPs (1.5,1.2, and 1.2-fold, respectively), but *not* with immune eQTLs excluded (0.9, 0.4, 0.5-folds, respectively). This suggests that enrichment of CNS SNPs in autoimmune traits was driven by overlapping immune eQTLs. We also observed enrichment of CNS SNPs for education years (1.25-fold), college completion status (1.29-fold), and BMI (1.55-fold), but neither waist-to-hip-adjusted BMI nor height showed enrichment of CNS SNPs.

**Figure 3.**
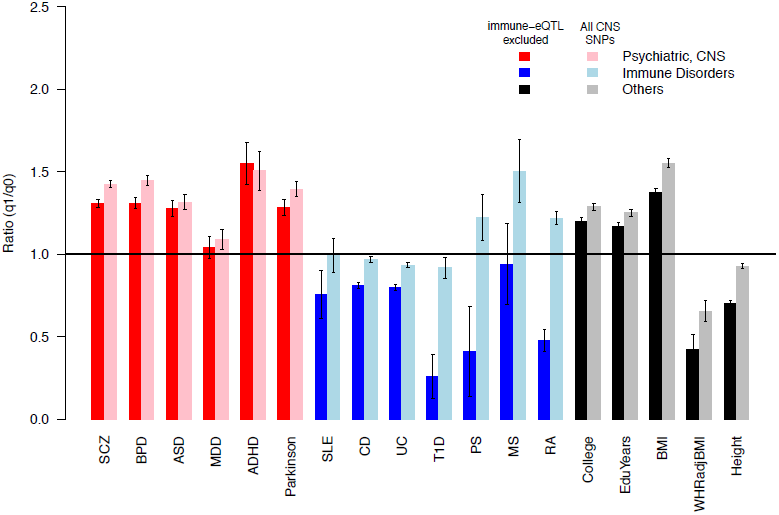
Enrichment of CNS SNPs in 18 traits. Enrichment of CNS SNPs (comprising 18.8% of all SNPs) in 18 traits from three categories: psychiatric disorders or CNS related disorder (red), immune system related disorders (blue), and body somatic features (black). For each trait, the first bar (darker color) excludes immune eQTLs from CNS SNPs, and the second bar (light color) is for all CNS SNPs (comprising 21.4% of all SNPs).

Next, we tested enrichment of immune eQTLs in the same set of 18 traits (**Fig. 4**). The seven immune-mediated disorders consistently had the strongest enrichment (ranging from 2.0 to 8.5-fold). We also observed enrichment of immune eQTLs in four psychiatric disorders (SCZ, BPD, ASD, and MDD; 2.0, 2.0, 1.4, 1.6-fold, respectively), and Parkinson’s disease (1.4-fold). Those enrichment effects still persisted with CNS SNPs removed, suggesting the enrichment was not solely due to overlapped CNS SNPs also being immune eQTLs. We also observed immune eQTL enrichment in two education traits, college completion (1.39-fold) and year of education (1.46-fold), and in three physical features, BMI (1.99-fold), obesity measured by waist-to-hip ratio adjusted BMI (2.90-fold), and height (2.97-fold).

**Figure 4.**
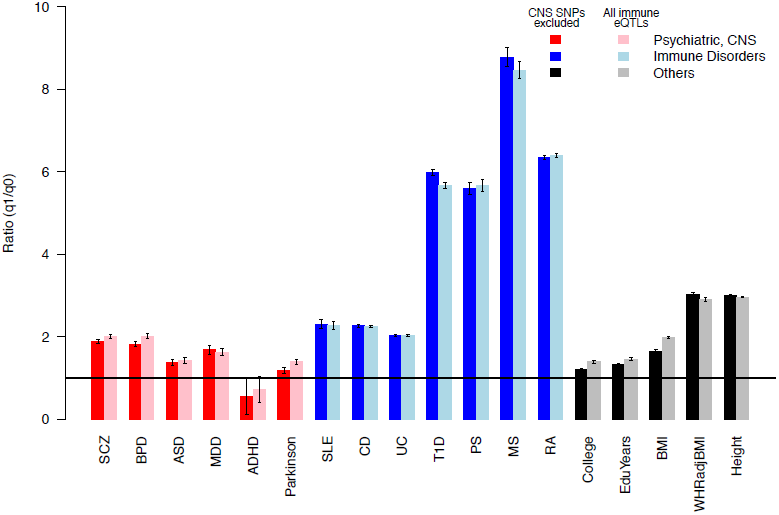
Enrichment of immune eQTLs in 18 traits. Enrichment of immune eQTLs (comprising 7.5% of all SNPs) in 18 traits from three categories: psychiatric disorders or CNS related disorder (red), immune system related disorders (blue), and other body somatic features (black). For each trait, the first bar (darker color) excluded CNS SNPs from immune eQTLs, and the second bar (light color) is for all immune eQTLs (comprising 10.1% of all SNPs).

There was even stronger enrichment of immune eQTLs among SNPs shared between five psychiatric traits (SCZ, BPD, ASD, MDD, ADHD) and CD (**Fig. S3** and **SI**). Next we tested enrichment of DNase-peak located SNPs in SCZ GWAS signals from 98 ENCODE tissues (**Table S7**), and found the top cell lines were from blood elements having important roles in immune response, with the top two cell lines being CD20+ B cells and Th2 cells (CD4+ T cells)(**Fig. S4**).

### Trend of consistent effect direction between psychiatric disorders and immune system disorders

To explore the mechanism of these pleiotropic effects further, we examined effect directions for SNPs having high posterior probabilities of being associated with both disorders. For each SNP, the same allele may increase or reduce susceptibility for the two disorders (same direction) or have opposite effects (different directions). The SCZ QQ plot (**Fig. S6**) shows a rise of the curve conditional on having the same effect direction with CD. We then considered four disorder pairs showing strong pleiotropy: SCZ-CD (*p*=1.9e-109), BPD-CD (*p*=1.5e-13), SCZ-Height (*p*=2.0e-122), and BPD-Height (*p*=5.4e-150). For each of these four pairs, there is no correlation in effect direction when *all* genotyped SNPs are considered (**Table S3**). This is expected because most SNPs are not associated with either trait. However, trends emerged after we partitioned the SNPs according to their posterior probabilities of being associated with both traits into 10 groups, and calculated the proportions of SNPs having the same effect directions for each group. There are clear patterns for SCZ-CD and BPD-CD, but not for SCZ-height nor BPD-height as shown in **Fig. 5**.

**Figure 5.**
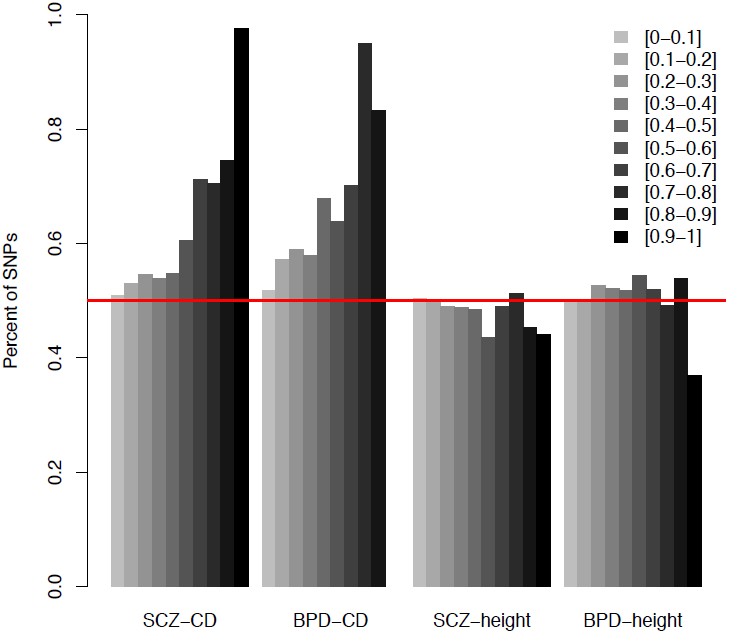
Trend of consistent effect directions for SCZ/BPD-CD across posterior probability groups. Proportion of SNPs having the same effect direction for trait pairs, in each of the 10 posterior probability groups (darker colors indicate higher posterior probability), where SNPs were grouped based on posterior of being associated with both traits into 10 equal bins. Four pairs of traits: SCZ-CD, SCZ-height, BPD-CD, and BPD-height.

**Figure 6.**
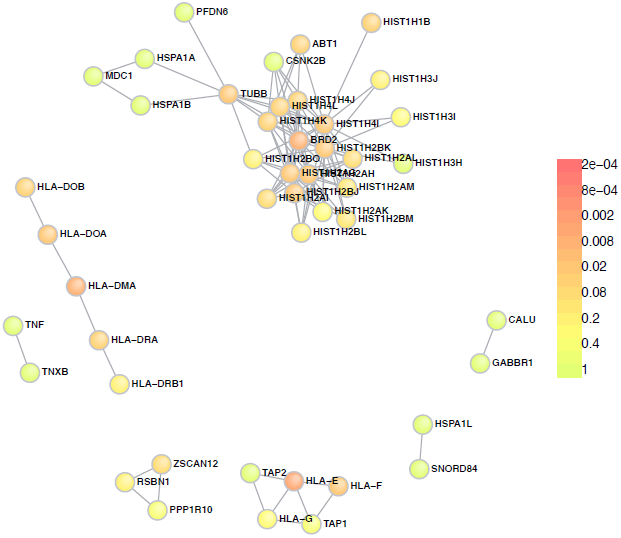
Protein-protein interaction enrichment. Constructed using the top 1000 SNPs, with color indicating significance level.

In general, the higher the posterior probability of a SNP being associated with SCZ (or BPD) and CD, the more likely that the SNP had the same effect direction for the pair. For SCZ-CD, among the 85 top SNPs with posterior probabilities of association with both SCZ and CD higher than 0.9 (**Table S4**), 97.6% of SNPs had the same effect directions (an allele either increases or reduces both SCZ and CD risks), with the only two opposite-direction SNPs near gene GALNT3. Similarly, for BPD-CD, in the SNP group with posterior probabilities higher than 0.8, 83% of SNPs had the same effect direction, and in the SNP group with posterior probabilities between 0.7 and 0.8, 95% of SNPs had the same effect direction for two disorders (**Fig. 5**). Similar patterns were also observed for SCZ-RA and BPD-RA pairs (**Fig. S7**). In contrast, for the SCZ-height pair, among the top SNPs with posterior probabilities of association with both SCZ and height higher than 0.9, 44% of the height-increasing alleles were SCZ risk alleles and the other 56% were SCZ protective alleles. In fact, the proportion of alleles that were associated with lower height and increased SCZ risk simultaneously was ∼50% for all SNP groups, regardless of their posterior probabilities of being associated with both SCZ and height. Effect direction distributions across 10 posterior groups for the BPD-height pair behaved similarly.

### Genome region enrichment analysis

To identify molecular pathways and networks underlying the connections between psychiatric disorders and immune-mediated disorders, we performed a series of genome region enrichment analysis. For each of the 28 disorder pairs (four psychiatric disorders and seven autoimmune disorders, with ADHD excluded due to its weak GWAS signals), we calculated the posterior probability of each SNP being associated with both traits, and plotted the posterior probabilities against genome positions (**Fig. S8**). A consistent peak in the chromosome 6 MHC region was observed for most trait pairs. There were also multiple additional recurrent or sporadic regions. To specify these regions and genes, next we performed cytoband enrichment analysis, protein-protein interaction (PPI) subnetwork detection, and gene ontology (GO) enrichment analysis.

Cytoband enrichments of SNPs with posterior probability > 0.5 were evaluated for each of the 28 disorder pairs using Fisher’s exact test. **Table S2** shows enriched cytobands with odds ratio (OR) > 5 and Bonferroni-adjusted *p* < 0.001. Under these stringent cutoffs, we found that the MHC region (6p22.1-21.3) was shared by psychiatric and autoimmune disorders (23 out of 28 pairs, **SI**). Apart from the MHC region, we also identified other shared regions (**Table S2**). For example, 1p13.2 was significantly enriched for the eight disorder pairs between [SCZ, BPD, MDD, and ASD] and [T1D and RA], with Bonferroni-adjusted p-values ranging from 6.8e-26 to 2.7e-78, with top SNPs located in genes *AP4B1, PTPN22,* and *PHTF1* (**Fig S9**).

With the hypothesis that there might be a common mechanism underlying the shared genetic liability between different psychiatric disorder-autoimmune disorder pairs in general, we selected SNPs with high confidence of affecting both classes of disorders (**SI**) and performed enrichment analysis for Gene Ontology (35) (GO) terms, KEGG pathways, and PPI networks. PPI can provide independent information for prioritization of genetic findings (36-39), and thus we constructed PPI sub-networks via DAPPLE (40) in which PPI edges are overrepresented in top SNPs (SNPs with posterior > 0.8 in at least 6 disorder pairs, **Fig. S5**). The sub-networks (**Fig. 7**) highlighted include the *HLA-D* subfamily; a transcription activation related sub-network including multiple histones, *BRD2, TUBB* and *ABT1*; and *HLA-E, HLA-F, TAP1* and *TAP2.* Genome annotation enrichment was performed via DAVID (41, 42) on GO (35) terms and KEGG pathways (43, 44), with gene lists constructed with genes containing SNPs having posterior > 0.8 in at least three disorder pairs (**Fig. S5**). The identified top terms of enrichment included “antigen processing and presentation,” “MHC protein complex,” “allograft rejection,” and “NF-kappaB binding” (**Table S5**), which further suggests the enrichment of immune system function in shared genetic factors between psychiatric and autoimmune disorders.

## Discussion

Our results demonstrate extensive shared genetic factors between psychiatric and autoimmune disorders. First, 24 of 35 psychiatric disorder-autoimmune disorder pairs we considered yielded significant evidence for pleiotropy based on GPA analysis of summary statistics from GWAS results. Second, enrichment of immune related eQTLs in multiple psychiatric disorder GWAS signals was observed, including SCZ, BPD, ASD, and MDD. Third, much higher enrichment of immune related eQTLs in SNPs shared by psychiatric disorders and CD was observed. Fourth, enrichment of psychiatric disorder GWAS signal was observed in DNase I hypersensitivity sites in cell lines based on cellular elements with important roles in immune response. Fifth, there was a clear trend of shared SNPs having the same effect direction for SCZ-CD, SCZ-RA, BPD-CD, and BPD-RA. Finally, analyses based on posterior probabilities of SNPs being associated with both psychiatric and autoimmune disorders yielded reoccurring genomic regions, with significantly enriched cytobands and pathways. All these different lines of evidence indicate that psychiatric disorders are rather strongly genetically related to autoimmune disorders.

Beyond the evidence of pleiotropy, our results suggest *how* psychiatric disorders and autoimmune disorders are related genetically. Consistent with previous research, we observed a major role of MHC region for both classes of disorders, which repeatedly stood out among multiple analyses. Specific proteins in MHC regions were prioritized via constructing overrepresented protein-protein interaction sub-networks. Specifically, those protein-protein interaction clusters most responsible for shared genetic components between psychiatric disorders and autoimmune disorders in these data were: (1) three minor gene subunits *HLA-E*, *HLA-F*, and *HLA-G*, but not the three major gene subunits, interacting with *TAP1, TAP2. TAP1* and *TAP2* are transporters associated with antigen processing, which cooperate with MHC class I to present antigens (45); (2) Interaction between HLA DO, DM, and DR proteins; and (3) a set of genes with important roles in transcriptional activation, including *BRD2, TUBB, ABT1,* and multiple histone coding genes.

The MHC is not the only genomic region contributing to the connection between psychiatric disorders and autoimmune disorders. First, we observed enrichment of immune eQTLs even after the whole MHC region was removed (**Fig. S10**). Second, cytoband enrichment results indicate roles played by other specific genomic regions, such as 1p13.2, harboring gene *PTPN22* (“Protein Tyrosine Phosphatase, Non-Receptor Type 22 (Lymphoid)”), which was also prioritized in our PPI analysis.

Our work pinpoints these specific genetic regions, genes, proteins, and pathways, which potentially connect psychiatric disorders and immune disorders genetically and therefore etiologically. These specific genes might be considered as starting points for follow-up experiments concerning the etiological linkage of those two classes of disorders. Furthermore, our findings might be used in developing novel drugs that treat psychiatric disorders by modulating immune system function, which would nicely complement the limited existing treatment options for these psychiatric disorders.

The observation of a tendency of the same effect direction for SNPs associated with either SCZ and BPD paired with CD gives some insight concerning the underlying mechanism of their shared genetic factors. Pleiotropy has been extensively reviewed (16, 46-48), but is still not well understood in terms of its extent, mechanisms, and consequences. The Weak Hypothesis of Universal Pleiotropy (WHUP) advocated by Fisher (49) and Wright (50) is based on two assumptions that, in general, a phenotype might be influenced by many variants, and a variant might cause changes to many phenotypes. This basic idea finds considerable support in GWAS findings, and fits the Common-Disease Common-Variant (CDCV) hypothesis. Under WHUP, extensive pleiotropy should be detected between phenotypes. In this case though, the effect directions of shared genetic variants should be about random. Therefore, our results suggest that it is unlikely that the shared genetic factors between psychiatric and autoimmune disorders originate from universal pleiotropy (or a tendency of GWAS to identify SNPs that are functional and for SNPs that are functional to be those that are most likely to influence various traits); rather, our observation supports a closer genetic relationship between those two types of disorders. Various molecular mechanisms could result in pleiotropy (48). There are biological pleiotropy, mediated pleiotropy, and spurious pleiotropy. Biological pleiotropy has separate causal paths for different phenotypes, while mediated pleiotropy has one phenotype lying on another phenotype’s causal path; thus by this mechanism, one phenotype might lead to another (48). Our results, the striking trend of shared SNPs for SCZ and CD acting in the same direction, can be best explained by mediated pleiotropy. This, together with our observation of pervasive enrichment of immune eQTLs in psychiatric disorders, and the lack of enrichment of CNS SNPs (immune eQTLs excluded) in immune-mediated disorders (except MS, which is characterized by CNS pathology), suggest that autoimmune disorders might mediate psychiatric disorder risk, i.e. some downstream autoimmune dysfunctions might be a trigger to some psychiatric disorders (or subtypes).

We observed considerable enrichment of immune-related eQTLs in traits of body features (height, BMI, and WHR-adjusted BMI), which are consistent with previous experiments that BMI is correlated with immune parameters (51), and that height is associated with immune response in young men (52). Therefore, our results further confirm the relationship between body somatic features (BMI and height) and immune system from a genomics perspective. The enrichment of CNS SNPs in two educational traits is consistent with the notion that educational attainment is related to CNS development.

Although our analyses were based on results from GWAS consortia, the statistical power remains limited to identify the majority of disease associated variants for these disorders. GWAS results from larger studies and improved statistical and bioinformatics approaches will enable us to identify more shared genetic pathways between these classes of disorders, and as always – despite the very high significance levels we observed for some relationships – independent replication of our results is called for.

## Materials and Methods

### Genome-wide Association Study (GWAS) Data Sources

We made use of GWAS summary statistics from a set of diverse and representative traits, including major psychiatric disorders, various immune system disorders, multiple metabolic diseases and metabolism related traits, body morphological features, and some socioeconomic measures (**Table 1**). The *p*-values were available for all traits, but only some of them have available specified alleles and their corresponding beta or odds ratios indicating effect direction.

### Genomic Annotation Data Sources

Central nervous system (CNS) genes were identified in a previous study (53), comprising preferentially brain-expressed genes (53), neuronal-activity genes (54), learning-related genes (55), and synapse genes, defined by Gene Ontology (35). A complete list of these genes is given in **Table S6**. CNS SNPs were defined as SNPs located within CNS genes with 50kb extension at each end. To investigate immune system influence, we used context-specific eQTLs upon triggering immune response as detected by Fairfax et al. (56), where interferon-γ and lipopolysaccharide (LPS) were used as inflammatory proxies to stimulate innate immune effects in monocytes from volunteers of European ancestry. We used a union of eQTLs detected in different contexts as a set of immune-related eQTLs in our study. In total, we have 94,674 immune eQTLs and 199,202 CNS SNPs, of which 24,860 CNS SNPs are also immune eQTLs. To investigate the impact of chromatin state, we used DNase I hypersensitivity sites extracted from ENCODE (57) DNase-seq peaks and signal of open chromatin from 125 cell lines.

### Statistical Methods

Pleiotropy analysis was performed via the GPA R package (31), which uses a statistical approach to exploring the genetic architecture of complex traits by integrating pleiotropy and functional annotation information, including prioritizing risk genetic variants, evaluating annotation enrichment and pleiotropy by hypothesis testing. GPA does not need individual level genotype-phenotype data, making it particularly useful for largescale integrative analysis. We briefly describe the parameters and their interpretations in the GPA model in **Supporting Information**, and more details can be found in (31).

Genome annotation enrichment was carried out using DAVID (41, 42); PPI sub-networks were generated using DAPPLE(40). Cytoband position information was downloaded from the UCSC Table Browser(58). Cytoband enrichment was tested using Fisher’s exact test, with *p*-values adjusted using Bonferroni correction Dunn (59).

## Supporting Information

### GPA method description

GPA (1) is a statistical approach to exploring the genetic architecture of complex traits by integrating pleiotropy and functional annotation information, including prioritizing risk genetic variants, evaluating annotation enrichment and pleiotropy by hypothesis testing. Instead of relying on genotype-phenotype data at the individual level, it only requires the summary statistics from GWAS, which makes it useful for integrative analysis of genomic data. For completeness, we briefly introduce the GPA model here.

Consider the *p*-values {*p*_1_..., *p*_*M*_} obtained by performing hypothesis testing of genomewide SNPs from one GWAS, where *M* is the number of SNPs. In the GPA model, these *p*-values are assumed to come from a mixture of null (un-associated) and non-null (associated), with probability *π*_0_ and *π*_1_ = 1 – *π*_0_, respectively. GPA uses the Uniform distribution on [0,1] and the Beta distribution with parameters (*α*, 1) to model the *p*-values from the null and non-null groups, respectively. Let *Z*_*j*_ ϵ {0,1} be the latent variable indicating whether the *j*-th SNP is from the null or non-null group, where *Z*_*j*_ = 0 means null and *Z*_*j*_ = 1 means non-null. Then the GPA model for one GWAS without annotation can be written as:

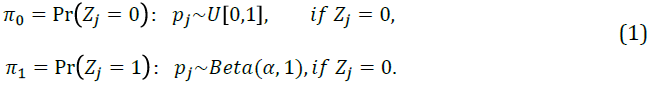

GPA further incorporates functional annotation as follows. Let an M-dimensional vector *A* collect functional information from an annotation source, where *A*_*j*_ ϵ {0,1} indicates whether the *j*-th SNP is a functional unit according to the annotation source. For example, given an eQTL data, if the *j*-th SNP is an eQTL, then *A*_*j*_ = 1, otherwise *A*_*j*_ = 0. The relationship between *Z*, and *A*_*j*_ is described as:

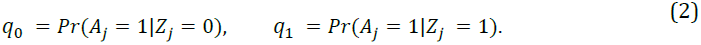

Clearly, *q*_0_ can be interpreted as the proportion of null SNPs being annotated, *q*_*1*_ corresponds to the proportion of non-null SNPs being annotated, and *q*_*1*_ > *q*_0_ implies that there exists enrichment in this annotation. In our study, the ratio *q*_*1*_/*q*_*0*_ is define as enrichment fold measuring the strength of enrichment in various annotations.

An efficient Expectation-Maximization (EM) algorithm has been developed to adaptively estimate the model parameters {*π*_0_, *π*_1_, *q*_0_, *q*_1_ *α*). After that, SNPs can be prioritized based on their local false discovery rates (FDR). When there is no annotation data, the local FDR is defined as the probability that the *j*-th SNP belongs to the null group given its *p*-value, i.e., *fdr*(*p*_*j*_) = *Pr*(*Z*_*j*_ = 0|*P*_*j*_). With annotation data, the FDR can be calculated as *fdr*(*p*_*j*_ , *A*_*j*_) = *Pr*(*Z*_*j*_ = 01, *A*_*j*_). We can use the likelihood ratio test to assess the significance of its enrichment. Specifically, the significance of enrichment of an annotation for GWAS can be assessed by testing *H*_0_ : *q*_0_ = *q*_1_ versus *H*_1_ : *q*_0_ ≠ *q*_1_. Standard errors of all the parameters can also be calculated.

The extension of the above model to handle two GWAS is straightforward. Suppose the *p*-values from two GWAS have been collected in an *M* × 2 matrix **p** = [*p*_*jk*_], where *p*_*jk*_ denotes the *p*-value of the *j*-th SNP in the *k*-th GWAS, *k* = 1, 2. Let *Z*_*j*_ 6 {00,10,01,11} indicate the association between the *j*-th SNP and the two phenotypes: *Z*_*j*_ = 00 means the *j*-th SNP is associated with neither of them, *Z*_*j*_ = 10 means it is only associated with the first one, *Z*_*j*_ = 01 means it is only associated with the second one, and *Z*_*j*_ = 11 means it is associated with both. Then the two-groups model (1) can be extended to the following four-groups model:

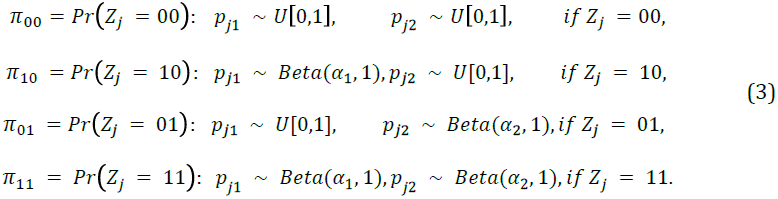

Similarly, functional annotation information can be incorporated into the multiple GWAS model (3) in the following way:

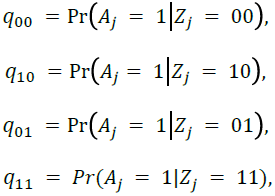

where *q*_00_ is the probability of a null SNP being annotated, *q*_10_ is the probability of the first phenotype associated-SNP being annotated, *q*_01_ is the probability of the second phenotype associated-SNP being annotated, and *q*_11_ is the probability of jointly associated-SNP being annotated. For joint analysis of two GWAS data sets, the local FDR calculation and enrichment assessment can be done in a similar way. In addition, the pleiotropy between two phenotypes can be tested in a statistically rigorous way. When there is no pleiotropy, i.e., the signals from the two GWAS are independent of each other, testing pleiotropy can be formulated by testing the following hypothesis:

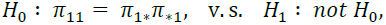

where *σ*_1*_ = *σ*_10_ + *σ*_11_ and *σ*_*1_ = *σ*_01_ + *σ*_11_. The likelihood ratio test statistic asymptotically follows χ^2^ distribution with *df* = 1 under the null.

More details about the GPA approach have been discussed in (1).

### Enrichment of immune-eQTLs in SNPs shared by psychiatric disorders and autoimmune disorders

To explore this hypothesis further, we tested levels of enrichment of immune related eQTLs in SNPs associated with both psychiatric disorders and Crohn’s disease, and observed large enrichment ratios. Our result shows that, among the immune related eQTLs, the ratios (*q*_11_/*q*_00_) of the group of SNPs associated with both diseases and the group of SNPs associated with neither diseases are SCZ-CD 3.9 (s.e.=0.06), BPD-CD 4.4(s.e.=0.09), ASD-CD 3.4(s.e.=0.2), MDD-CD 4.6(s.e.=0.18), and ADHD-CD 2.1(s.e.=0.45) (**Fig. S3**). The ratios with respect to SNPs associated with only one disease (*q*_10_/*q*_00_ and *q*_01_/*q*_00_) are much lower, suggesting that the shared genetic components between the 5 psychiatric disorders and CD are closely related to immune function.

### Enrichment of DNase-Peak SNPs in schizophrenia GWAS

DNase-Peak dataset was downloaded from ENCODE for 125 cell lines, and there were 98 cell lines after removing 27 cancer cell lines (**Table S7**). Although limited in cell lines from brain regions, those 98 cell lines have great coverage for various blood cells, making it suitable for studying whether there is enrichment of functional genomic regions in SCZ GWAS tissues implicated with important immune functions. DNase-Peak SNPs are SNPs located in or within 1kb from DNase-Peaks. A cluster of blood originated cell lines is the top cell lines in our analysis (**Fig. S4**).

### Cytoband enrichment test based on posterior probability

Cytoband position was downloaded from the UCSC Table Browser(2), with 862 entries of cytobands in total. Enrichment tests were carried out on 28 pairs of disease pairs, between seven autoimmune disorders (CD, UC, MS, PS, RA, SLE, and T1D) and four psychiatric disorders (SCZ, BPD, MDD, and ASD). For each disease pair, potential shared SNPs were selected based on posterior of being associated with both diseases Pr(*Z*_*y*_ = 1)>0.5. Numbers of potential shared SNPs vary from disease to disease, ranging from 0 to 4,505 (for SCZ-CD). For each cytoband, we calculated {*X*_11_, *X*_10_, *X*_01_, *X*_00_}, with *X*_11_ being the number SNPs in cytoband that are potential shared SNPs, *X*_10_ being the number of SNPs in cytoband that are not potential shared SNPs, *X*_01_ being the number of potential shared SNPs not in cytoband, and *X*_00_ being the number of SNPs not in cytoband and are not potential shared SNPs. We then tested the deviation from null hypothesis that (*X*_11_, *X*_10_, *X*_01_, *X*_00_} follows hypergeometic distribution using Fisher’s exact test and adjust *p*-values for multiple testing using Bonferroni correction Dunn (3).

A complete list of all cytobands with enrichment odds ratio (OR) >5 and Bonferroni-adjusted *p*-value<0.001 in at least one disease pair were reported (**Table S2**). Some cytobands have significant enrichment in more than one disease pairs, such as MHC region and 1p13.2, indicating their role in affecting both psychiatric disorders and autoimmune disorders.

For example, we observed very strong evidence of MHC region implicated in affecting both classes of disorders. In eight psychiatric disorder-IBD pairs (between SCZ, BPD, MDD, and ASD; and UC and CD), four pairs have significant enrichment for 6p22.1 and 6p21.32, and 3 pairs for 6p21.33. In 20 psychiatric disorder-autoimmune disorder pairs (between SCZ, BPD, MDD, and ASD; and MS, PS, RA, T1D, and SLE), 19 pairs have significant (*OR*>5 and Bonferroni adjusted *p*-value<0.001) enrichment for 6p22.1, 6p21.33, 6p21.32, eight pairs have significant enrichment for 6p21.31, and 6 pairs for 6p22.2.

### Supplementary Figures

**Figure S1.**
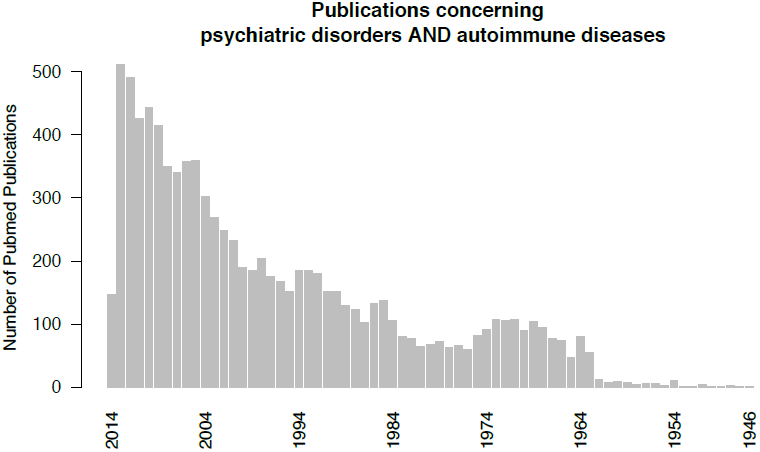
Number of publications from PubMed search of “Psychiatric disorders AND autoimmune diseases” until Oct. 2014 per year.

**Figure S2.**
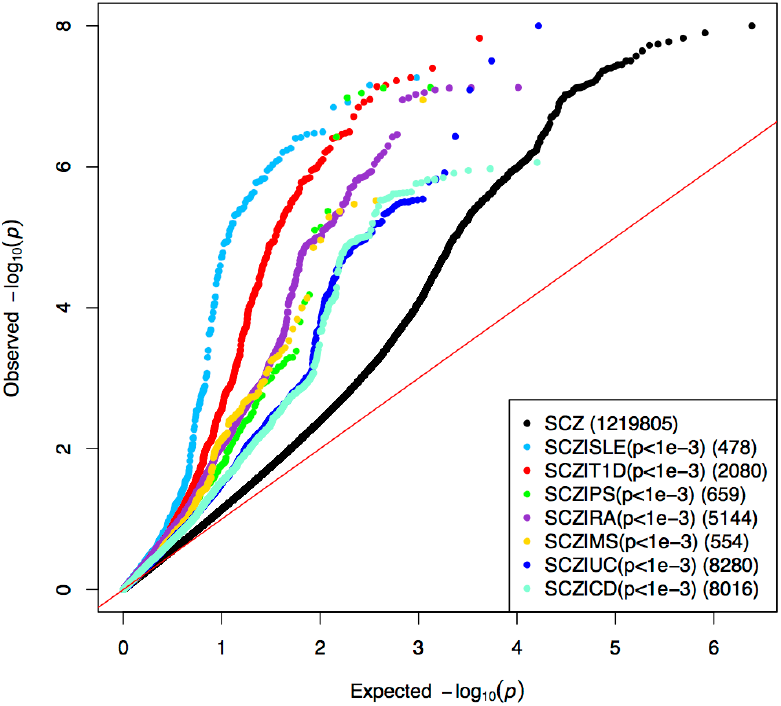
QQ plot showing pervasive pleiotropic effects between SCZ and seven immune mediated disorders. Black dots represent all 1,219,805 SCZ GWAS SNPs while the other 7 colored dots represent different subsets of SNPs selected from the corresponding autoimmune disorder GWAS whose p values were < 0.0001, with the number of SNPs in each subset shown in brackets.

**Figure S3.**
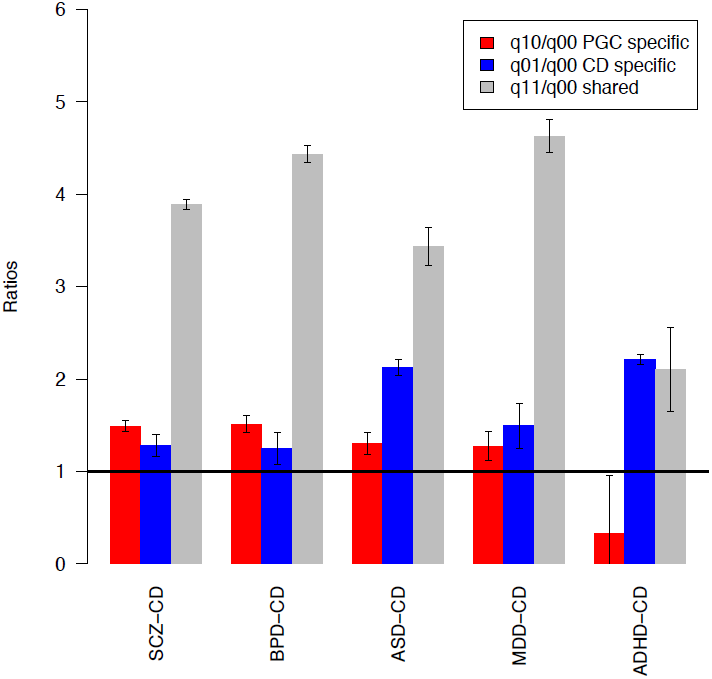
Enrichment of immune-eQTLs in SNPs shared between five psychiatric disorders and CD. Red bars represent the enrichment ratio of immune related eQTLs in 5 PGC traits specific SNPs respectively, blue bars represent the enrichment ratio of immune related eQTLs in SNPs associated only with CD, grey bars shows the enrichment ratio of immune related eQTLs in SNPs associated with both CD and the corresponding PGC disorder.

**Figure S4.**
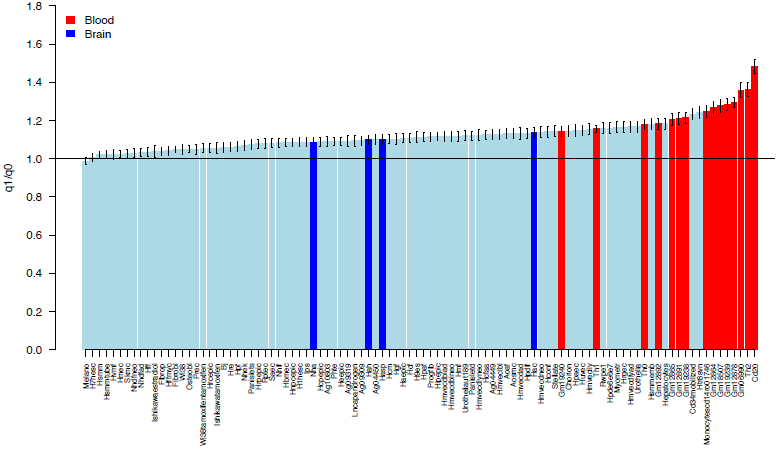
Enrichment of DNase-peak located SNPs in SCZ GWAS signal from 98 ENCODE cell lines. 98 cell lines ordered by enrichment ratios; cell lines from blood and brain are colored red and blue, respectively.

**Fig S5.**
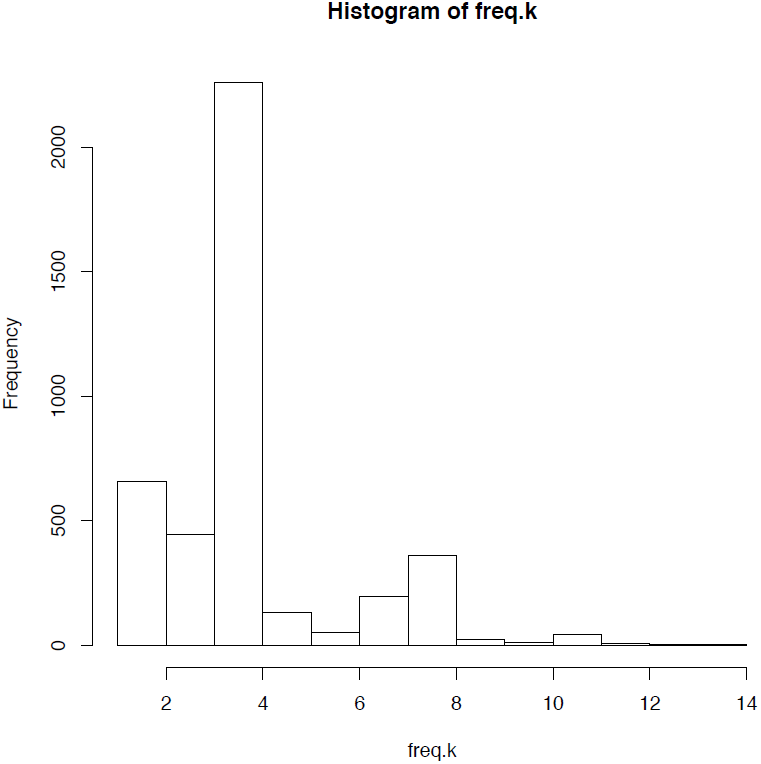
The frequencies of top SNPs (posterior>0.8) appearing in 35 disease pairs. A total of 4,149 SNPs have posterior probability>0.8 of being associated with at least one disease pair. Histogram showing the number of disease pairs that these 4,149 SNPs are associated with (with posterior probability>0.8).

**Figure S6.**
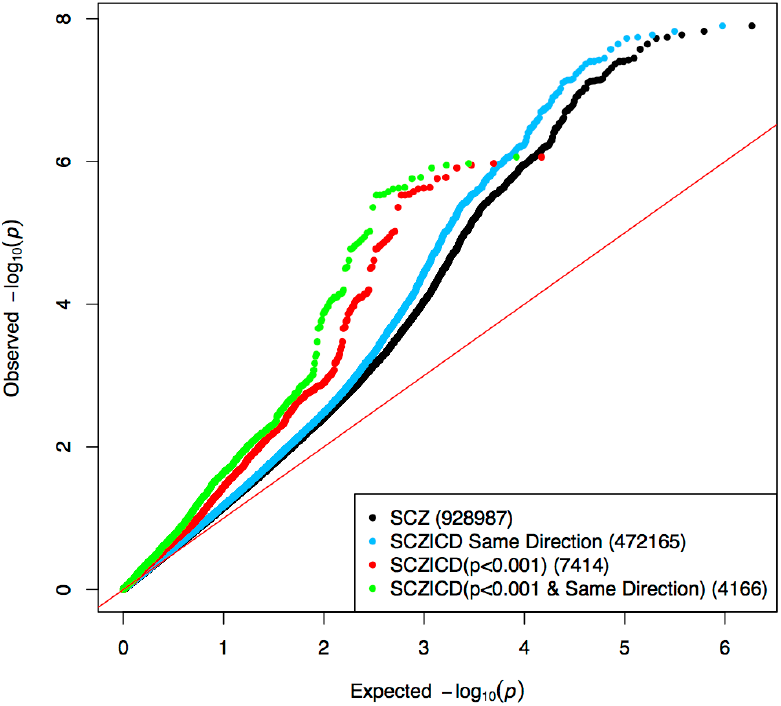
QQ plot showing enrichment of SNPs having same effect direction between SCZ and CD. Black dots for all 928.987 SNPs, and the other three lines are different subsets of SNPs, selected by: blue for 472,165 SNPs that have same effect direction for SCZ and CD; red for 7,414 SNPs with p<0.001 in CD GWAS; green for the intersection of blue and red.

**Figure S7.**
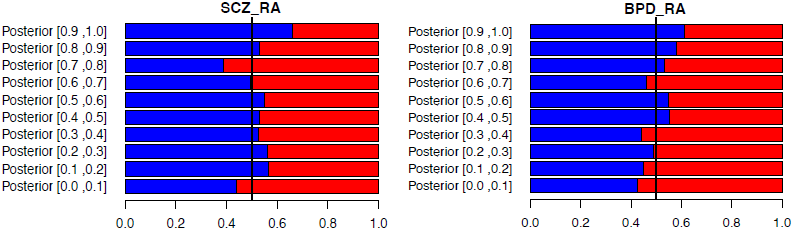
Trend of consistent effect directions for shared SNPs of SCZ-RA and BPD-RA disease pairs. For each disease pair, SNPs are assigned to 10 groups based on their posterior probability of being associated with both diseases, and then the proportion of alleles having the same effect direction for the disease pair was calculated within each of the ten SNP groups. Left, SCZ-RA; right, BPD-RA. Blue represents the same effect direction, and red represents the opposite direction, x-axis represents proportion of SNPs.

**Figure S8.**
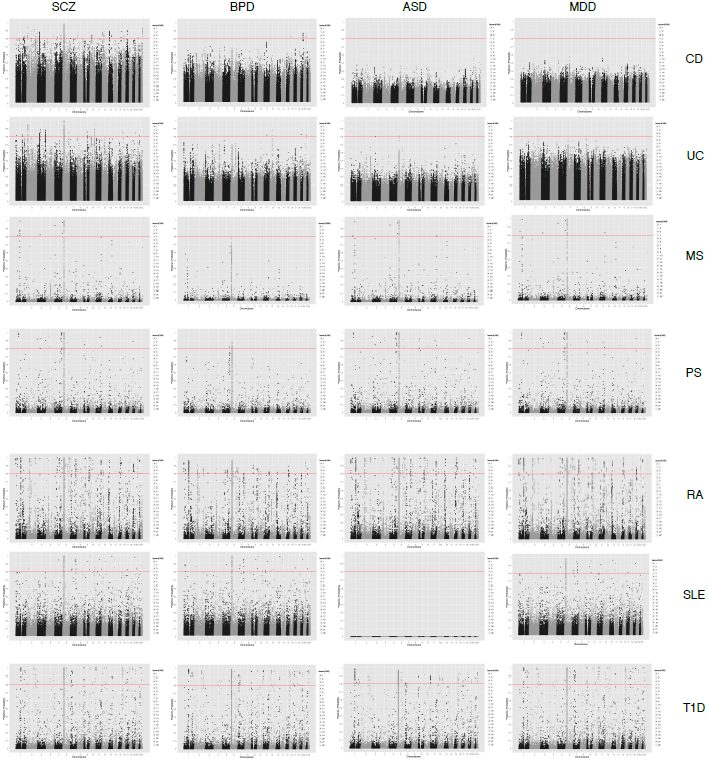
Posterior of SNPs being associated with both diseases for 28 disease pairs. Posterior of all shared SNPs of a disease pair is plotted against 22 genomic positions. Red line indicates posterior=0.8.

**Figure S9.**
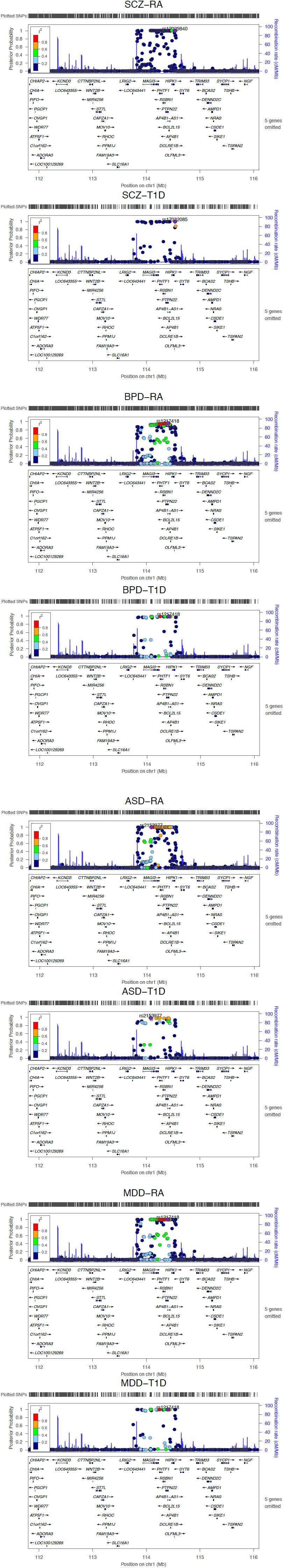
LocusZoom showing distribution of posterior probability in cytoband 1p13.2. Posterior probability of being associated with both diseases in eight disease pairs between SCZ, BPD, MDD, ASD, and RA, T1D, are shown separately. Only region 1p13.2 is shown. SNP with the highest posterior is labeled, and SNPs in LD with it are colored. Figures plotted using LocusZoom (4).

**Figure S10.**
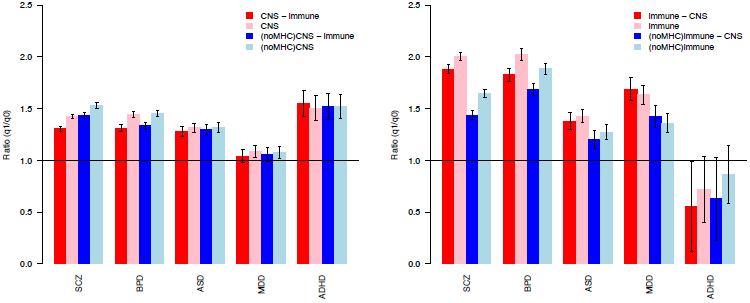
Enrichment of CNS SNPs and immune eQTL with and without MHC region. Red bars are enrichment ratios estimated using all SNPs genome-wide, blue bars are enrichment ratios estimated with SNPs in MHC region excluded.

Table S1 (Separate Excel File)

Strength of pleiotropy within and across two disorder classes: 5 psychiatric disorders and 7 autoimmune disorders.

Table S2 (Separate Excel File)

Cross-disorder cytoband enrichment results.

**Table S3.**
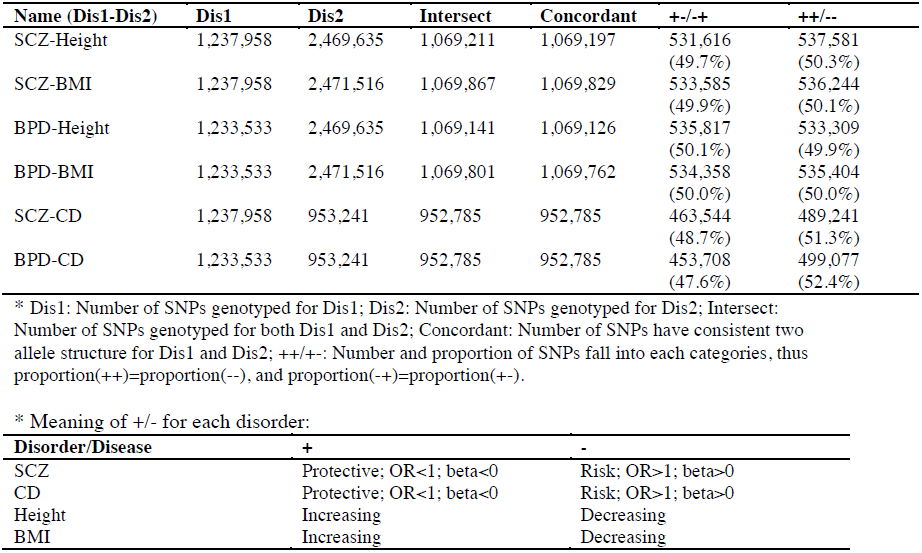
Number of SNPs in analysis and proportions of SNPs in direction combination categories

Table S4 (Separate Excel File)

A list of 85 SNPs with posterior probability of being associated with both SCZ and CD
above 0.9.

**Table S5.** Genome Annotation Enrichment Results (adjusted *p*-value<0.05)

| Category | Term | Fold.Enrichment | Bonferroni |
| --- | --- | --- | --- |
| KEGG_PATHWAY | Allograft rejection | 16 | 5.46E-09 |
| GOTERM_BP_FAT | antigen processing and presentation | 11 | 1.16E-08 |
| GOTERM_CC_FAT | MHC protein complex | 14 | 2.28E-08 |
| KEGG_PATHWAY | Type I diabetes mellitus | 14 | 3.52E-08 |
| GOTERM_MF_FAT | NF-kappaB binding | 20 | 2.61E-07 |
| KEGG_PATHWAY | Autoimmune thyroid disease | 12 | 3.37E-07 |
| GOTERM_CC_FAT | flotillin complex | 48 | 3.59E-07 |
| KEGG_PATHWAY | Systemic lupus erythematosus | 7 | 7.40E-07 |
| KEGG_PATHWAY | Antigen processing and presentation | 8 | 7.41E-07 |
| GOTERM_BP_FAT | antigen processing and presentation of peptide antigen | 21 | 7.76E-07 |
| GOTERM_CC_FAT | nucleosome | 12 | 1.18E-06 |
| GOTERM_BP_FAT | response to unfolded protein | 11 | 3.39E-06 |
| GOTERM_MF_FAT | thyroid hormone receptor activator activity | 37 | 3.40E-06 |
| GOTERM_MF_FAT | thyroid hormone receptor coactivator activity | 37 | 3.40E-06 |
| KEGG_PATHWAY | Graft-versus-host disease | 13 | 5.22E-06 |
| GOTERM_BP_FAT | defense response | 3 | 6.10E-06 |
| GOTERM_CC_FAT | chromatin | 6 | 6.46E-06 |
| KEGG_PATHWAY | Viral myocarditis | 8 | 1.29E-05 |
| KEGG_PATHWAY | Cell adhesion molecules (CAMs) | 6 | 3.08E-05 |
| GOTERM_CC_FAT | protein-DNA complex | 9 | 3.39E-05 |
| GOTERM_BP_FAT | response to protein stimulus | 8 | 4.57E-05 |
| GOTERM_MF_FAT | retinoid-X receptor activity | 39 | 5.68E-05 |
| GOTERM_CC_FAT | MHC class I protein complex | 18 | 6.19E-05 |
| GOTERM_MF_FAT | receptor activator activity | 24 | 8.36E-05 |
| GOTERM_BP_FAT | nucleosome assembly | 8 | 0.000247 |
| GOTERM_MF_FAT | retinoic acid receptor activity | 29 | 0.000343 |
| GOTERM_BP_FAT | antigen processing and presentation of peptide antigen via MHC class I | 24 | 0.000351 |
| GOTERM_BP_FAT | chromatin assembly | 8 | 0.000356 |
| GOTERM_BP_FAT | protein-DNA complex assembly | 8 | 0.000567 |
| GOTERM_CC_FAT | TAP complex | 44 | 0.000603 |
| GOTERM_MF_FAT | transcription factor binding | 3 | 0.000705 |
| GOTERM_BP_FAT | nucleosome organization | 8 | 0.000709 |
| GOTERM_CC_FAT | plasma membrane | 2 | 0.00072 |
| GOTERM_BP_FAT | immune response | 3 | 0.00101 |
| GOTERM_CC_FAT | chromosomal part | 3 | 0.00117 |
| KEGG_PATHWAY | Asthma | 12 | 0.00196 |
| GOTERM_CC_FAT | MHC class I peptide loading complex | 34 | 0.00211 |
| GOTERM_BP_FAT | chromatin assembly or disassembly | 6 | 0.00246 |
| GOTERM_MF_FAT | MHC class I receptor activity | 21 | 0.00249 |
| GOTERM_MF_FAT | retinoic acid receptor binding | 21 | 0.00249 |
| GOTERM_MF_FAT | receptor regulator activity | 14 | 0.00333 |
| GOTERM_MF_FAT | transcription regulator activity | 2 | 0.00393 |
| GOTERM_MF_FAT | protein heterodimerization activity | 4 | 0.00438 |
| GOTERM_BP_FAT | positive regulation of immune system process | 4 | 0.00453 |
| GOTERM_MF_FAT | MHC class II receptor activity | 19 | 0.00456 |
| GOTERM_MF_FAT | advanced glycation end-product receptor activity | 59 | 0.00675 |
| GOTERM_BP_FAT | DNA packaging | 6 | 0.00704 |
| GOTERM_CC_FAT | plasma membrane part | 2 | 0.00809 |
| GOTERM_MF_FAT | ligand-dependent nuclear receptor transcription coactivator activity | 11 | 0.01 |
| GOTERM_MF_FAT | vitamin D receptor binding | 15 | 0.0124 |
| GOTERM_CC_FAT | chromosome | 3 | 0.0147 |
| GOTERM_BP_FAT | response to nutrient | 5 | 0.0385 |
| GOTERM_MF_FAT | DNA binding | 2 | 0.0389 |
| KEGG_PATHWAY | Intestinal immune network for IgA production | 7 | 0.0408 |
| GOTERM_MF_FAT | hormone receptor binding | 6 | 0.042 |
| GOTERM_MF_FAT | MHC class I protein binding | 18 | 0.0434 |
| GOTERM_CC_FAT | caveola | 8 | 0.0477 |

Table S6 (Separate Excel File)

A complete list of central nervous system (CNS) genes.

Table S7 (Separate Excel File)

A list of description of 98 ENCODE cell lines.

